# Coevolutionary maintenance of forked tails and song in hirundines (Aves: Hirundininae)

**DOI:** 10.1101/2022.09.25.509392

**Authors:** Masaru Hasegawa

## Abstract

Both conspicuous plumage ornamentation and song are well-known examples of sexually selected traits but their interrelationship is not well-known, perhaps in part because of confounding factors, including interspecific variation in ecology, habitat, morphology, and type of ornamentation. Here, using a phylogenetic comparative approach, we examined the evolutionary relationship between forked tails and the presence/absence of song in hirundines (Aves: Hirundininae). Hirundines have similar ecology (e.g., aerial insectivores, social monogamy, and biparental provisioning), morphology (e.g., syrinx with nearly complete bronchial rings), and plumage ornamentation (i.e., a sexually selected forked tail), which provides a unique opportunity to examine the evolutionary associations between plumage ornamentation and song. In particular, hirundines have repeatedly lost their ornamentation, forked tails, enabling us to test their association with the evolutionary gain/loss of their simple song. After controlling for phylogeny and covariates, we demonstrated that song was less likely to be found in species with forkless tails than in species with forked tails. Two correlates of tail shape, sexual dimorphism in the overall plumage characteristics as a well-known measure of sexual selection and incubation type as a measure of extrapair mating opportunity, had no detectable relationship with the presence/absence of song, indicating the importance of forked tails, rather than their correlates. Evolutionary pathway analysis further supported the correlated evolution of the two traits, in which forked tails and song are maintained together and less likely to be lost under the presence of each other. The current study provided macroevolutionary support for the integrated use of visual and acoustic courtship traits.

---

Sexual selection explains the evolution of conspicuous traits, even if they are costly for survival (Andersson 1994). Such conspicuous “ornamental” traits are found across modalities, including several visual signals (e.g., conspicuous coloration/patterning and exaggerated body appendages) and acoustic signals (e.g., melodious song), even in the same clade (e.g., Aves). In fact, both intra- and interspecific analyses support sexual selection as an explanation for the evolution and maintenance of each of these traits (e.g., see reviews in Andersson 1994; Hill & McGraw 2006; Tazzyman et al. 2014; Rosenthal 2017).

The evolutionary relationships among these conspicuous traits across modalities, however, remain unclear. A classic “transfer hypothesis” predicts that visual ornamentation and songs are negatively correlated across bird species (i.e., evolutionary tradeoff between them), because the resources invested in a costly signal cannot be used for investing in another, leading an evolutionary tradeoff between the two costly signals (Darwin 1971). On the other hand, positive correlations among cross-modal signals can also be expected, for example, when they provide compensatory information (e.g., van Doorn & Weissing 2004) or are functionally integrated (as “courtship phenotype”; Ligon et al. 2018). In fact, some comparative studies have revealed positive associations between them (e.g., Shutler & Weatherhead 1990; Gonzalez-Voyer et al. 2013; Ligon et al. 2018), others revealed negative associations (e.g., Badyaev et al. 2002; Laverde et al. 2017; Beco et al. 2021; Tietz et al. 2022), and the remaining studies found no detectable associations (e.g., Ornelas et al. 2008; Soma & Garamszegi 2015; Matysiokova et al. 2017). Possible explanations for the lack of consistency among (and even within) studies include the confounding effects of interspecific variation in habitats, mating systems, condition, and ornament type (e.g., Shutler & Weatherhead 1990; Badyaev et al. 2002; Schutler 2011; Mason et al. 2014; Ligon et al. 2018).

Swallows and martins (Aves: Hirundininae) offer a unique opportunity to test the evolutionary association between plumage ornamentation and song. First, they have similar ecological niches, such as aerial insectivores, highly adapted to aerial locomotion, typically socially monogamous, biparental provisioning species (Turner & Rose 1994), and thus the confounding effects of variation in ecology are at best small. Second, hirundines often have sexually selected “forked” tails (e.g., Møller 1988, 1989; reviewed in Møller 1994; Turner 2006; Romano et al. 2017), but otherwise lost their forked tails in the evolutionary time scale and instead have forkless tails (e.g., Turner & Rose 1994; Hasegawa & Arai 2022). Repeated evolutionary loss of forked tails provides a rare opportunity to test evolutionary associations among the conspicuous traits across modalities. In other words, we can test whether loss of visual signal has been associated with the evolution of acoustic signal using this clade. Evolutionary loss of forked tails is explained by reduced sexual selection, but not by foraging mode nor changing direction of sexual selection (Hasegawa & Arai 2022), supporting the importance of sexual selection for the evolution and maintenance of forked tails. Third, Hirundininae has a syrinx (i.e., a vocal organ of birds) having more or less complete bronchial rings, which differs from the syrinx having half bronchial rings found in other passerine birds (and in the other hirundine subfamily, Pseudochelidoninae: Turner & Rose 1994), suggesting that their song evolution might differ from other passerines because of this unique syrinx structure. In fact, hirundines sing relatively simple songs, and others lack apparent songs (and utter only “calls”, i.e., vocalizations much simpler than songs; Turner & Rose 1994). This is particularly important, because previous studies often focused on birds with complex songs (e.g., finches; Badyaev et al. 2002; Soma & Garamszegi 2015), and thus they focused on the elaboration process of a given signal (e.g., variation of complex song), rather than the initial evolutionary stages (e.g., gain/loss) of song. Here, we focus on the initial evolutionary stages of song, which might differ from the subsequent elaboration process (e.g., exaptation and potential sexual conflict for signal honesty matter in the subsequent elaboration process; Hill 1994; Bergstrom & Dugatkin 2016). Furthermore, an analysis of complex, elaborated song is difficult to evaluate (e.g., Mason et al. 2014), and different axes sometimes show conflicting results (e.g., some song axes provide support for a positive association whereas other axes provide support for a negative association even in the same study; e.g., Shutler & Weatherhead 1990; Ornelas et al. 2008; Beco et al. 2021). The presence/absence of song would not suffer from this difficulty (i.e., the presence/absence of song can be regarded as the presence/absence of conspicuous vocalization).

Using a phylogenetic comparative analysis, we here examined the evolutionary relationship between tail shape (forkless/forked) and the presence/absence of song in hirundines. We predicted that hirundines with forked tails would be more likely to have song than hirundines with forkless tails, because sexual selection complementarily or synergistically favors visual and acoustic signals, at least in the barn swallow *Hirundo rustica*, a model species for studying sexual selection (e.g., Møller et al. 1998; Wilkins et al. 2015), and thus the evolutionary loss of one composite would result in the evolutionary loss of another composite of the courtship phenotype. To account for the potential confounding effects of sexual dimorphism in the overall plumage characteristics and extrapair mating opportunity, both of which are positively correlated with tail shape (Hasegawa & Arai 2022), we also examined these measures in relation to the presence/absence of song. Lastly, we examined evolutionary transitions among states with and without forked tails in relation to the presence/absence of song to test whether the two traits evolved and maintained together.

## Methods

### Data collection

As in our previous study (e.g., Hasegawa & Arai 2018, 2020a,b), information concerning morphological traits (i.e., wing length and tail shape) and migratory habits was obtained from Hasegawa & Arai (2020b), which was in turn obtained from Turner & Rose (1994). Information on tail shape can be extracted from Turner & Rose (1994), and all hirundine species have either forked or forkless (square) tails (i.e., no species have pintails, graduated or round tails: Hasegawa & Arai 2022). We focused on the evolutionary loss of ornamentation (i.e., forked to forkless tails; Hasegawa & Arai 2022) to clarify the evolutionary association between plumage ornamentation and song (see Introduction). For this reason, we did not adopt an alternative approach using the depth of forked tails as a continuous variable (and in fact including forkless tails as a 0-mm depth forked tails are problematic: Hasegawa & Arai 2022). Detailed information of data collection is given elsewhere (Hasegawa & Arai 2020b). In short, the mean wing length of the species was used as representative of species body size to account for the potentially confounding effect of body size, because wing length is tightly correlated with other body size measures (e.g., body mass) and did not impose a reduction of sample size. For wing length, whenever sex-specific values were available, values from males were used as before (e.g., Hasegawa & Arai 2018, 2020b). The migratory habits were divided into two categories: migrants and others. We regarded species as migrant when they had breeding sites separated from their wintering sites (i.e., when they were a summer visitor in a portion of their distribution range; Turner & Rose 1994).

We also collected data concerning the presence/absence of song based on the descriptions in the “voices” category of each species (i.e., whether or not “song” is noted in the species) in Turner & Rose (1994). Of course, this dichotomous classification is only a rough measure of song, because whether the focal species have “song” in addition to much simpler “calls,” is dependent on the human observers, but this measure would still capture some interspecific variation in vocal complexity (i.e., songs are considered much more complex than calls). Further differentiation of the songs (e.g., melodious or not) was not used because of limited information. Similarly, because of limited information, we could not differentiate song into further subcategories based on its functional usage (e.g., courtship song, territorial song; note that their functions are often overlapped; Turner 2006). We searched for updates using del Hoyo et al. (2020) and similar approaches based on the literature description were found in preceding studies (e.g., Webb et al. 20019). We did not use vocal recordings on internet platforms (e.g., xeno-canto), because such information is still limited and is often lack the detailed information at data collection (note that different from photograph or similar visual information, acoustic information is difficult to identify, particularly when prior information on song is lacking). Of 72 species, we obtained the presence/absence of song from 69 species (i.e., the remaining three species lack any information of vocalizations from the literature).

Finally, we focused on two correlates of tail shape (Hasegawa & Arai 2022), sexual dimorphism in overall plumage characteristics as a measure of sexual selection and extrapair mating opportunity, measured as incubation type (female-only or biparental incubation), to study its relationship with the presence/absence of song (Table S1; see Hasegawa & Arai 2022 for detailed information). In short, we regarded the focal species as sexually dimorphic when the literature noted sex differences in the plate (and otherwise, the focal species was regarded as sexually monomorphic). Concerning a proxy of extrapair mating opportunities (frequent/rare), we used incubation type (female-only versus biparental incubation) from a previous study (Hasegawa & Arai 2020b), which was in turn obtained mainly from Turner & Rose (1994). Incubation type was tightly linked to extrapair paternity both within and between species in this clade (Magrath & Elgar 1997; Hasegawa & Arai 2020b). Dataset can be found in Table S1.

### Phylogenetic comparative analysis

We used a Bayesian phylogenetic mixed model with a binomial error distribution to examine the presence/absence of song. As before (e.g., Hasegawa & Arai 2020a,b), to account for potentially confounding factors [body size, measured by log(wing length), and migratory habit], we included these variables along with sexual tail monomorphism/dimorphism in the model. To account for phylogenetic uncertainty, we fit the models to each tree and applied a multimodel inference using 1000 alternative trees for swallows from birdtree.org (e.g., Garamszegi & Mundry 2014; Hasegawa & Arai 2017), which is sufficient to control for phylogenetic uncertainty (Rubolini et al. 2015). The residual variance was fixed at 1, as recommended for categorical dependent variables (de Villemereuil et al. 2013). We derived the mean coefficients, 95% credible intervals (CIs) based on the highest posterior density, and MCMC-based P-values, together with the phylogenetic signal (de Villemereuil & Nakagawa 2014). Similarly, we also tested whether the presence/absence of song could be explained by each of the ecological correlates of forked tails (see the previous section). All analyses were conducted in R ver. 4.0.0 (R Core Team 2020) using the function “MCMCglmm” in the package “MCMCglmm” (ver. 2.29: Hadfield 2010). We ran the analysis for 140,000 iterations, with a burn-in period of 60,000 and a thinning interval of 80 for each tree. The reproducibility of the MCMC simulation was assessed by calculating the Brooks– Gelman–Rubin statistic (Rhat), which must be <1.2 for all parameters (Kass et al. 1998), by repeating the analyses three times.

We used a discrete module in BayesTraits (Pagel 1999) to examine evolutionary transitions among states with tail shape (forkless/forked) and the presence/absence of song. Again, we used 1000 trees to account for phylogenetic uncertainty (see above). Here, we ran for 1,010,000 iterations with a burn-in period of 10,000 and a thinning interval of 1000 (note that the phylogenetic tree was randomly chosen from 1000 trees in each iteration). We denoted means as the representatives of each transition rate. We also showed the posterior probability that the differences in transition rates would be higher (or lower) than zero (as P_MCMC_ values). The reproducibility of the MCMC simulation was assessed by calculating the Brooks– Gelman–Rubin statistic (Rhat), which must be <1.2 for all parameters (Kass et al. 1998), by repeating the analyses three times. Bayes factor was calculated by comparing the marginal likelihood of a dependent model that assumed the correlated evolution of sexual dimorphism and biparental incubation to that of an independent model that assumes the two traits evolved independently, using the stepping stone sampler implemented in BayesTraits. Bayes factors of >2 indicate positive evidence. As before (e.g., Hasegawa & Arai 2022), we confirmed that statistical results supporting correlated evolution (see “Results” section) represented replicated co-distributions and bursts throughout hirundines and thus false positives due to a few influential evolutionary events would be unlikely (Maddison & FitzJohn 2015; Revell & Hermon 2022).

## Results

### Inter-relationship between visual and acoustic traits

Of the 69 species with available information (i.e., 96% of all 72 species; see Methods section), 14 species had no apparent song; the remaining 55 species had songs. Ancestral character reconstruction indicates that song was lost multiple times in this clade (Fig. 1).

**Fig. 1.**
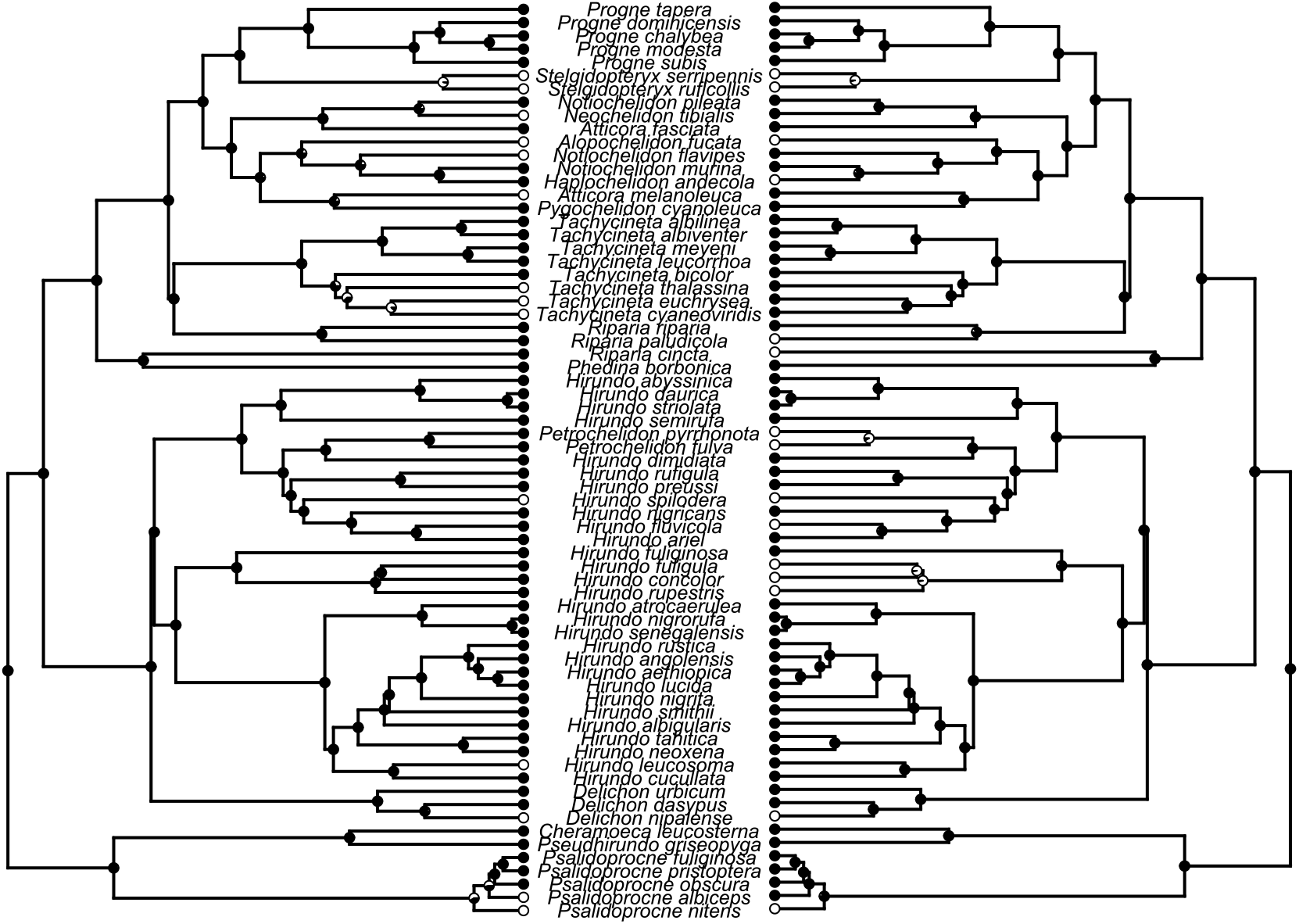
Examples of ancestral character reconstruction of the song (left) and of forked tails (right) in swallows and martins (Aves: Hirundininae). On the left side, the presence and absence of song are indicated with black and white circles, respectively. On the right side, species with forked tails and species with forkless tails are indicated with black and white circles, respectively. Likewise, the proportions of black and white at the nodes indicate the probability of the ancestral state.

The presence/absence of song was explained by tail shape: species with forkless tails were less likely to have song (Table 1b; Fig. 1). A correlate of tail shape, sexual plumage dimorphism (see Methods), was not significant when we used this variable instead of the presence/absence of song (Table 1). When we used extrapair mating opportunity measured as incubation type, another correlate of tail shape (see Methods), this variable was far from significant (Table 1), indicating that these well-known measures of sexual selection (i.e., sexual plumage dimorphism and extrapair mating opportunity) could not explain the observed positive relationship between tail shape and song.

**Table 1.** Multivariable Bayesian phylogenetic mixed model with a binary error distribution predicting the presence/absence of song in relation to (a) tail shape (n = 69), (b) sexual dimorphism in overall plumage characteristics (n = 69), and (c) extrapair mating opportunity measured by incubation type (n = 41) of swallows and martins (Aves: Hirundininae).

| Variables | Coefficient [95% CI] | $P_{MCMC}$ |
| --- | --- | --- |
| (a) Models with tail shape |  |  |
| Intercept | **-56.90 [-110.12, -6.04]** | **<0.01** |
| log(wing length) | **12.25 [0.60, 23.09]** | **<0.01** |
| Migratory habit | 0.26 [-1.85, 2.35] | 0.79 |
| Tail shape (forkless/forked) | **2.56 [0.40, 5.02]** | **0.02** |
| Mean phylogenetic signal = 0.75 |  |  |
| (b) Models with sexual plumage dimorphism |  |  |
| Intercept | **-55.92 [-108.56, -6.94]** | **0.01** |
| log(wing length) | **12.50 [1.66, 23.83]** | **0.01** |
| Migratory habit | 0.10 [-1.80, 2.04] | 0.93 |
| Sexual plumage dimorphism | -1.00 [-3.23, 1.07] | 0.37 |
| Mean phylogenetic signal = 0.72 |  |  |
| (c) Models with extrapair mating opportunity |  |  |
| Intercept | -38.16 [-103.86, 39.98] | 0.30 |
| log(wing length) | 8.41 [-8.27, 22.08] | 0.27 |
| Migratory habit | 0.94 [-1.57, 3.99] | 0.47 |
| Incubation type (female-only/biparental) | 0.24 [-2.66, 3.12] | 0.86 |
| Mean phylogenetic signal = 0.74 |  |  |

Model-averaged coefficients and 95% credible intervals (CI) are shown.
Bold indicates significant variables.

### Evolutionary transitions between states

When examining evolutionary transitions among states, we found that a dependent model that assumed a correlated evolution of tail shape and the presence/absence of song better fit the data compared to an independent model that assumes the two traits evolved independently (Bayes factor = 3.01, indicating strong evidence for correlated evolution). In the dependent model, the transition rate from the state with forked tails and song to the state with forked tails and no apparent song was lower than vice versa (P_MCMC_ = 0.024; Fig. 2). Also, the transition rate from the state with forked tails and song to the state with forkless tails and no song was lower than vice versa (P_MCMC_ = 0.016; Fig. 2).

**Fig. 2.**
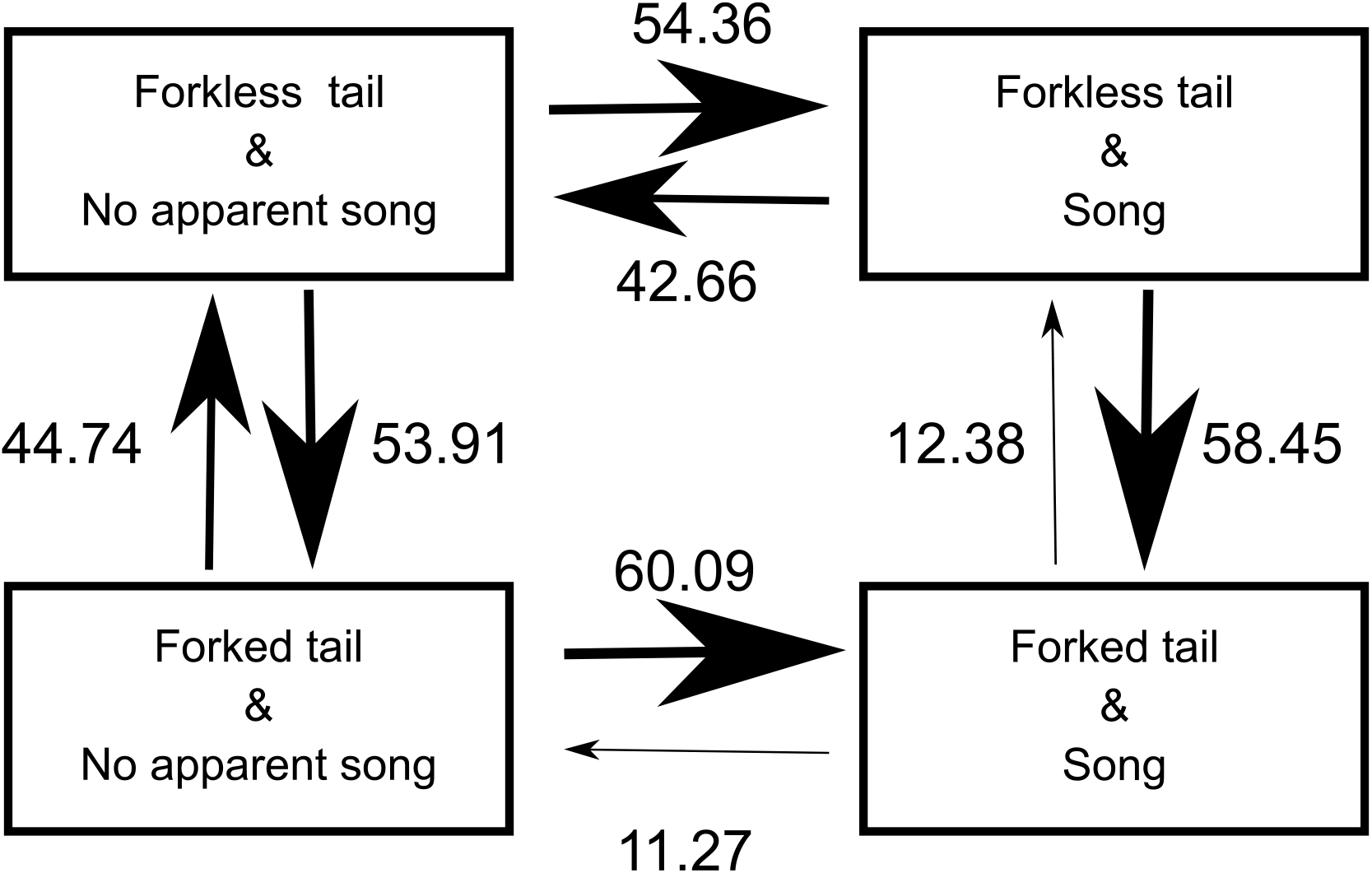
The most likely evolutionary transitions between states with different tail shape and the presence/absence of song in swallows and martins (Aves: Hirundininae). Numbers next to arrows, reflected by arrow size, indicate transition rates between pairs of states.

## Discussion

The main finding of the current study is the positive evolutionary association between forked tails and the presence of song. Evolutionary pathway analysis indicates that forked tails and song are maintained together and less likely to be lost under the presence of another trait. Because the presence/absence of song, our criterion here, is based on the literature description (see Methods), no apparent song does not mean they lack acoustic signals, and instead means a lack of complex vocalizations (note that every species has some sort of “calls” in the current dataset; see Methods). One may also think the possible observation bias (i.e., song might be overlooked in less-studied species). Although we could not completely exclude this possibility, this would not likely be the case, because there is in general a study bias towards studies in temperate region, i.e., breeding sites of migratory birds (Langmore 1998), but we did not find out the effect of migratory habit, suggesting that the possible confounding effect of observation bias would be at best small. Although further studies are clearly needed, we thus conclude that the observed positive relationship between sexual tail dimorphism and song is unlikely to be explained by observation bias alone (note that species with deeply forked tails, rather than species with forkless tails are more likely to have extinction risk, i.e., relatively “rare” species; Hasegawa & Arai 2018).

The observed positive relationship is intuitive, because the barn swallow, a model species for studying sexual selection, has both deeply forked tails and a “melodious” song (Turner & Rose 1994). Several empirical studies in fact show sexual selection for sequential or simultaneous evaluation of the two signals (e.g., Møller et al. 1998; Wilkins et al. 2015). As suggested by several researchers (e.g., courtship phenotype; Ligon et al. 2017; also see Wilkins et al. 2015 for the systems biology approach), our evolutionary pathway analysis supported the integrated use of cross-modal signals. The use of multi- modal signals might be adaptive (and thus be maintained) when it provides high joint benefits, such as multiple information (e.g., short-term condition via acoustic signal whereas long-term condition via plumage signals; Møller et al. 1998) or increased honesty as a whole (van Doorn & Weissing 2006), even if the joint cost of multiple signals might be high. Because song production, which requires a continuous change of vocal tracts (Riede et al. 2006), involves pulsating movement of body parts (including tails, i.e., the distal end of the body), the detectability of tail ornaments would also be enhanced by using song as a component of courtship phenotype (note that detectability of each trait is often assumed to be independent of each other e.g., Price & Schluter 1993). This might be particularly important to point out, because previous studies often ignore such byproduct effects of song. These are likely explanation at least in theory, but the actual “joint function” of forked tails and song during courtship remains to be clarified.

Although the abovementioned explanations are based on relatively “static” view of the interrelationship of traits, we should also stress that this might not always be applicable to any cross-modal signals. In fact, after the initial gain of signals, sexual conflict for honest signaling (or more broadly, coevolutionary feedback between signal senders and receivers) complicates the relationship between multiple candidate signals (Hill 1994; van Doorn & Weissing 2006), indicating the importance of changing benefits/costs of signals through its elaboration process. Thus, additional factor explaining why we could find out a positive evolutionary association is simply because we focused on the initial evolutionary stage of the courtship traits, in which the influence of sexual conflict might be small. Such evolutionary process-dependent associations between traits are likely, but most comparative studies to date unfortunately focused on already elaborated song and their further elaboration (see Introduction), and thus we lack the data to test this possibility. Future studies focusing on the initial evolutionary stage of song like ours (also see Webb et al. 2019 for female song), rather than subsequent elaboration, would be fruitful.

In summary, we found a positive evolutionary association between sexual tail dimorphism and the presence/absence of song, indicating a coevolutionary maintenance of the visual and acoustic signals. Because both positive and negative relationships were found between the visual and acoustic signals even in closely related taxa (e.g., different families of passerines; see Introduction), future studies should carefully take into account not only ecological factors but also evolutionary processes (i.e., initial evolutionary stage or subsequent elaboration process, here; see the preceding paragraph), both of which can affect the evolutionary associations of the signals.

## Acknowledgments

We thank Dr Emi Arai, Dr Shumpei Kitamura and his lab members at Ishikawa Prefectural University for their kindest advices. We are grateful to Dr Angela Turner for her kindly support on the valuable information on swallows. MH was supported by the Research Fellowship of the Japan Society for the Promotion of Science (JSPS, 22J40066).

---

**Table S1.**
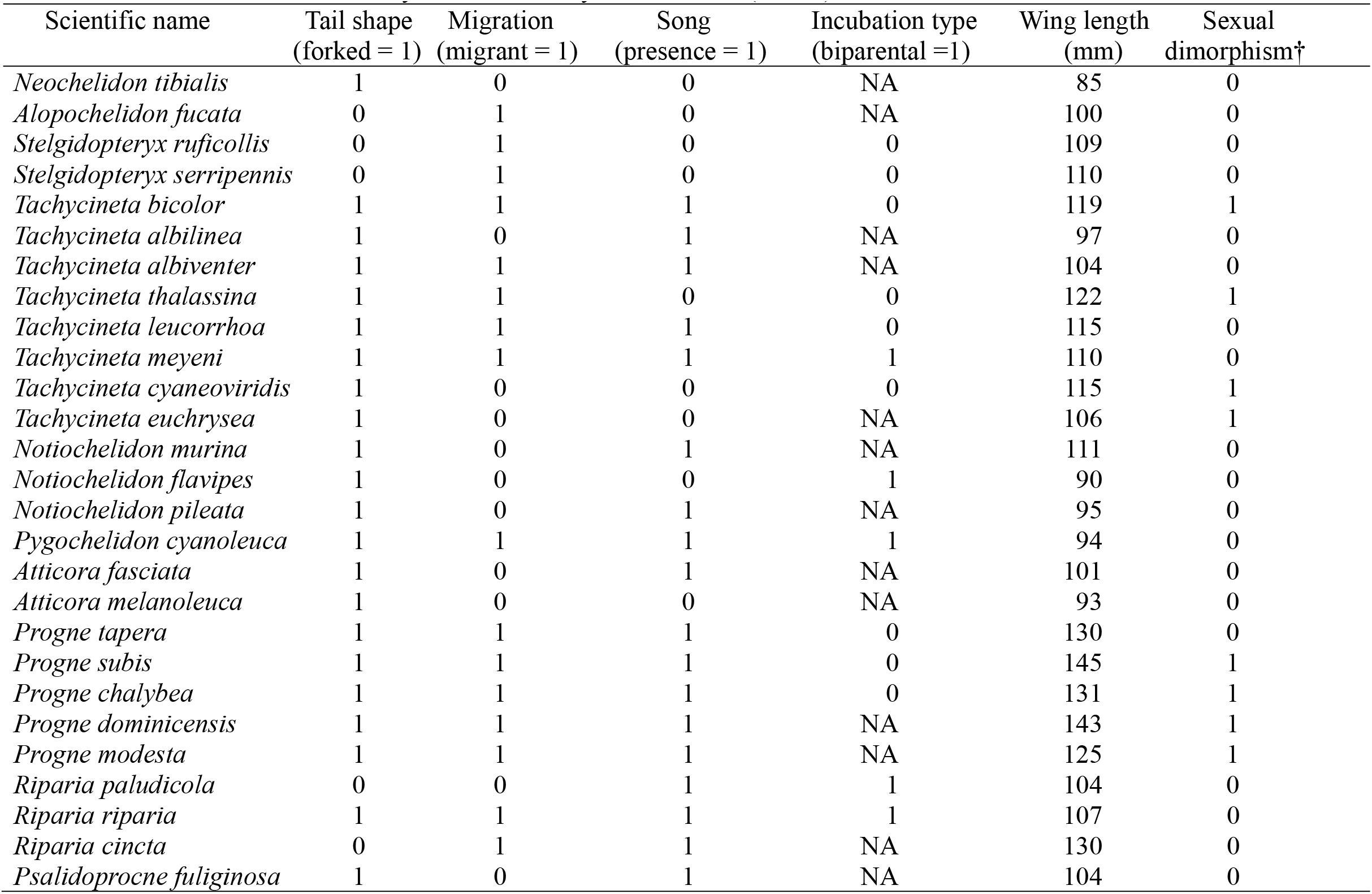
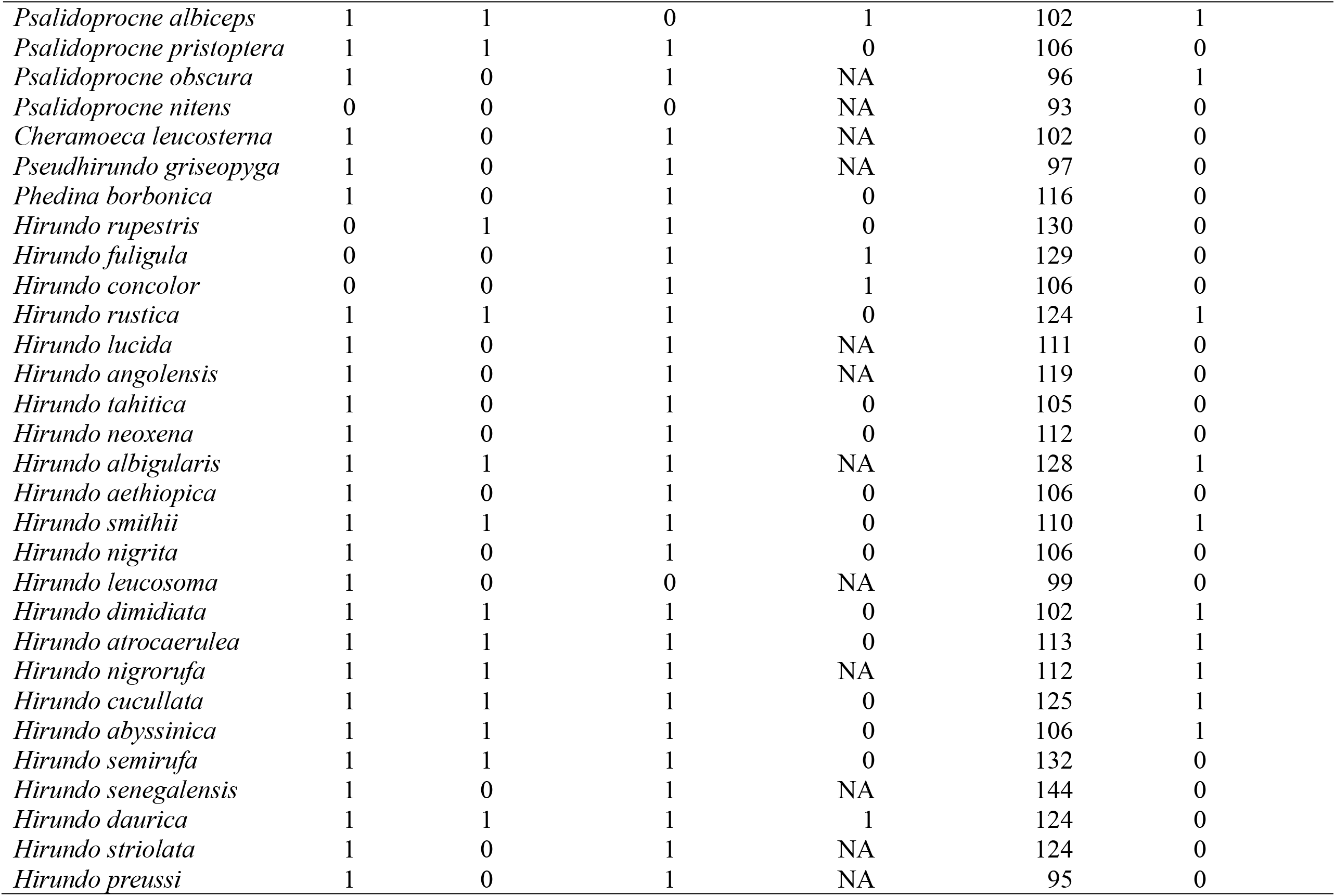
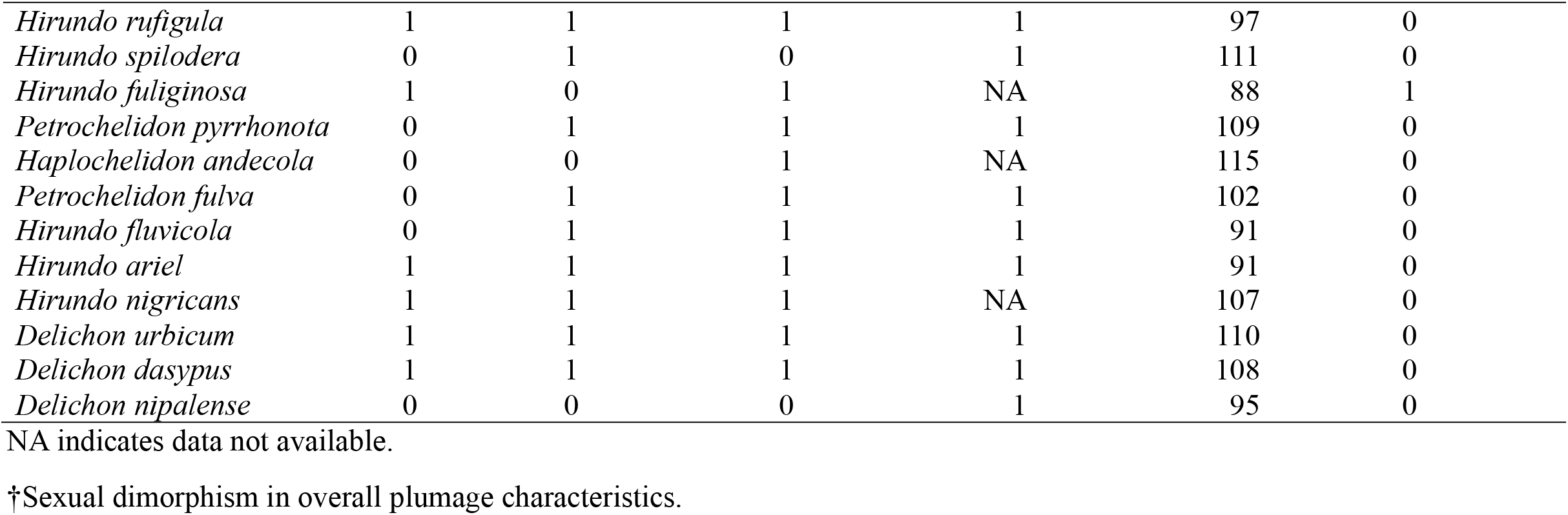
Data set for the current study in the subfamily Hirundininae (n = 69).

